# slideimp: Efficient Imputation for DNA Methylation Data

**DOI:** 10.64898/2025.12.15.694394

**Authors:** Hung Pham, Adam P. Lombroso, E. Cansu Cevik, Hugh S. Taylor, Kieran J. O’Donnell

## Abstract

**Motivation:** There is a growing need for efficient imputation methods for high-dimensional DNA methylation (DNAm) datasets. Existing microarray imputation approaches, such as k-nearest neighbors (K-NN) or principal component analysis (PCA)-based methods, provide high accuracy but can be computationally intensive, while methods for whole-genome data are not designed for large cohorts. We developed slideimp, an R package that implements sliding window, groupable, parallelized K-NN and optimized PCA imputation to address these limitations.

**Results:** Benchmarks on microarray DNAm datasets demonstrate that slideimp achieves up to 150x faster runtime and 10x-100x lower memory usage while maintaining comparable or superior accuracy over existing methods and implementations. For K-NN and PCA imputation, slideimp supports grouped imputation which enhances imputation efficiency and accuracy. In a whole-genome dataset, sliding window K-NN imputation substantially increased the correlation of the Horvath 2013 clock with chronological age from 0.131 to 0.477. Additional features include targeted imputation of CpG subsets for K-NN and estimation of imputation accuracy via repeated cross-validation. The efficient and flexible DNAm imputation methods implemented by slideimp can easily be applied to other high-dimensional data types.

**Availability and Implementation:** The code to fully reproduce all analyses presented in this paper is available on GitHub at https://github.com/hhp94/slideimp_paper. The R package slideimp is also available on GitHub at https://github.com/hhp94/slideimp. Supplemental figures are available online.

## Introduction

DNA methylation (DNAm), the addition of a methyl group to a cytosine nucleobase, typically cytosine-phosphate-guanine (CpG), is an abundant epigenetic modification in eukaryotic genomes and among the most studied epigenetic features in humans (Aristizabal *et al*., 2020). In large-scale epidemiological analyses, DNAm is typically profiled using DNA microarrays, such as the HumanMethylation450 (450K, ≈450,000 sites), MethylationEPIC v1.0 (EPICv1, ≈850,000 sites), or v2.0 (EPICv2, ≈930,000 sites) arrays from Illumina. The recently released Methylation Screening Array (MSA, Illumina) targets approximately 280,000 sites implicated in a variety of health conditions (Goldberg *et al*., 2025). Whole genome bisulfite sequencing (WGBS) or Enzymatic Methyl-seq (EM-seq) offer alternative approaches to describe variation in DNAm sites across the entire genome but at a higher per-sample cost.

Missing data is an important consideration for all DNAm analyses, especially for sequencing-based approaches where total missingness rate can exceed 30% at a standard 30x coverage. K-nearest neighbors (K-NN) imputation, a non-parametric imputation method from the impute package, is among the most widely used imputation methods for microarray DNAm data (Troyanskaya *et al*., 2001); while imputePCA (missMDA R package) is a principal component analysis (PCA)-based method that offers higher imputation accuracy at a greater computational cost (Josse and Husson, 2016). Other DNAm microarray imputation approaches include regression-based (methyLImp2) and random forest-based imputation (missRanger) (Plaksienko *et al*., 2024; Mayer, 2025). For whole-genome DNAm data, Hidden Markov Model-based imputation (METHimpute) and gradient boosting-based BoostMe are commonly used approaches (Taudt *et al*., 2018; Zou *et al*., 2018).

With the decreasing per-sample cost for both whole-genome and DNAm microarray analyses, there is a need for more computationally efficient and flexible imputation methods for extremely high-dimensional data. For example, for whole-genome DNAm imputation, BoostMe is no longer actively maintained and METHimpute does not make use of between-sample information to improve accuracy. In contrast, accurate imputation of DNAm microarrays can already be achieved using existing methods (e.g., impute.knn from the impute package), but efficiency could be improved by parallelization, which can significantly reduce runtime in large microarray DNAm datasets. Here, we introduce slideimp, a package that implements a more efficient K-NN imputation and optimized PCA imputation for DNAm data. To reduce runtime, slideimp’s K-NN imputation utilizes parallelization, adds support for tree-based nearest neighbor search strategies (i.e., KD-tree or Ball-tree) from the mlpack package, and allows users to impute all measured or targeted subsets of CpGs (Bentley, 1975; Omohundro, 1989; Curtin *et al*., 2023). Figure 1A shows a helpful decision tree to select the appropriate functions in slideimp.

**Figure 1.**
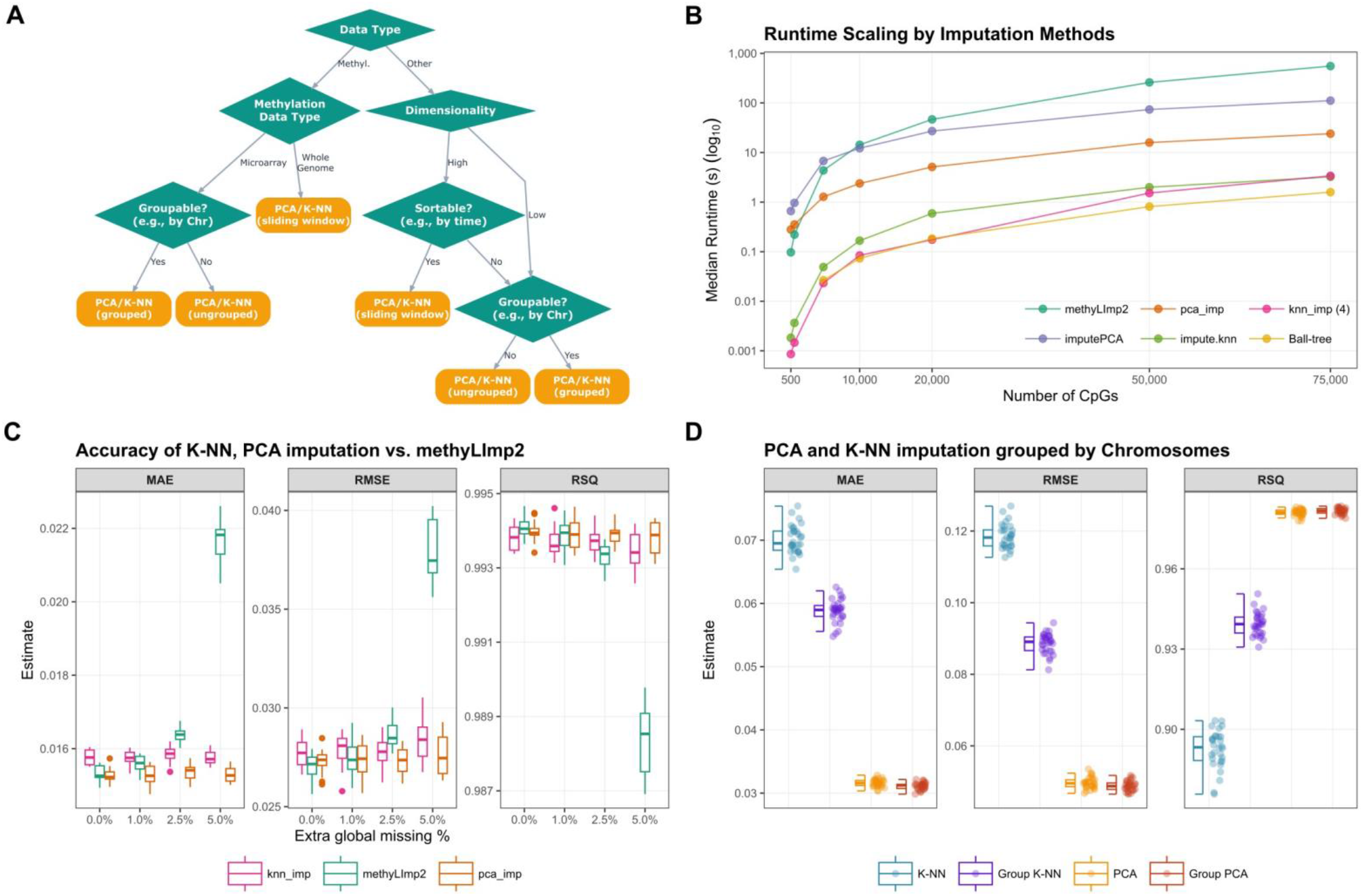
Overview of the slideimp package. (A) Decision tree for functions in slideimp: The package handles large DNA methylation data or other data types (e.g., microarrays or intensively sampled longitudinal data). Variables should be grouped before applying sliding window or normal imputation to reduce computational cost. slideimp also includes a helper function, tune_imp, to perform repeated cross-validation and measure the performance of K-NN vs. PCA, which aids in choosing the appropriate method. (B) Runtime benchmarks of different imputation methods: Points represent the median runtime measured in seconds. pca_imp (orange), knn_imp (4) (pink), and Ball-tree (yellow) represent the PCA, K-NN with 4 cores, and K-NN Ball-tree imputation implementations of slideimp, respectively. (C) Comparison of imputation accuracy between K-NN, PCA, and methyLImp2: With no added missingness (0.0%), all methods perform well for these data (GSE286313, EPICv1, N=72, blood samples, chromosome 22), with PCA imputation having the lowest MAE and methyLImp2 having the lowest RMSE and highest RSQ. methyLImp2 accuracy decreases with increasing missingness. (D) Comparison of imputation accuracy between K-NN and PCA grouped by chromosome vs. ungrouped: PCA imputation is significantly more accurate than K-NN imputation using these data (GSE264438, MSA, N=581, diverse tissues). The grouped imputation is faster and more accurate for both methods. Chr: chromosome; PCA: principal component analysis; K-NN: k-nearest neighbors; CpG: cytosine–phosphate–guanine site; MAE: Mean Absolute Error; RMSE: Root Mean Square Error; RSQ: R^2^.

We showcase slideimp using real-world data and simulations, highlighting improvements in runtime, memory usage, and imputation accuracy as compared to current best practices for DNAm data imputation. In addition, we highlight how slideimp’s sliding window K-NN or PCA imputation can be used to improve the accuracy of epigenetic age estimates from established biomarkers of aging (so-called ‘epigenetic clocks’), among the most widely used tools in epigenetic epidemiology, using both microarray and whole-genome-based experiments.

## Methods

### Data

For DNAm microarrays, we used data publicly available from NCBI GEO: the GSE286313 data (N=72) which consists of blood samples from four cohorts of mixed sex and diverse geographical origin, spanning early to late adulthood, with DNAm profiled using both the EPICv1 and EPICv2 arrays (Zhuang *et al*., 2025) and the GSE264438 data (N=581) which measures DNAm from various human cell lines and tissues on the MSA microarray (Goldberg *et al*., 2025). For whole-genome DNAm data, we used EM-seq data from blood samples of 41 reproductive-age women (mean age of 32.9 years, SD of 7.1) without pregnancy or malignancy recruited from the Yale New Haven Hospital from 2023 to 2025.

### Benchmarks of slideimp’s K-NN and PCA imputation

First, we benchmarked runtime and memory usage of slideimp’s K-NN and PCA imputation implementations using the GSE286313 data (EPICv1, chromosome 1). We compared slideimp imputation methods against impute.knn, the original imputePCA, and methyLImp2. For impute.knn, we configured a single two-means clustering step when N CpGs ≥1,500 for comparable accuracy. Each measurement was taken at least five times.

Next, we compared slideimp’s K-NN and PCA imputation accuracy against methyLImp2 using the GSE286313 data (chromosome 22) with increasing total missing rates from 0% to 5%. We calculated the mean absolute error (MAE), root mean squared error (RMSE), and R^2^ (RSQ) of the imputed values using repeated cross-validations with 14 repeats.

We then characterized slideimp’s feature to perform K-NN or PCA imputation with grouping, i.e., by chromosome. Using the MSA GSE264438 data (full microarray), for 30 replications, 300 CpG sites were randomly chosen, and 20 values per CpG were randomly set to missing. Imputations were performed using K-NN and PCA either grouped by chromosomes or ungrouped and the MAE, RMSE, and RSQ of the imputed values were compared.

Finally, we used Monte Carlo simulation with 1,000 iterations to evaluate epigenetic clock regression analyses with K-NN or PCA imputation under increasing per-CpG missingness (20% to 70%) with different missing data mechanisms. Data can be missing completely at random (MCAR), missing at random (MAR), or missing not at random (MNAR) (Rubin, 1976). For illustrative purposes we focused on chromosome 1 from the GSE264438 data. We simulated clock values from 500 chosen CpGs. Then, for each simulation iteration, we generated an outcome Y that is associated with these clock values with a β-coefficient of 0.1 and a signal-to-noise ratio of 0.1. We then generated missing data under the MCAR, MAR, MNAR mechanisms with different missing rates using the package missMethods, imputed the data, and regressed Y on the imputed clocks (M. S. Santos *et al*., 2019). We evaluated the performance of mean imputation, K-NN imputation, PCA imputation, and Ball-tree K-NN imputation by comparing the median and 95% quantile interval of the estimated imputed β-coefficient and standard error to the known true values.

### Epigenetic clocks estimation in whole-genome DNAm data

Whole blood EM-seq DNAm data from 41 reproductive age women with no pregnancy or malignancy were processed with the methylKit package (Akalin *et al*., 2012). Missing values were imputed by 1) a baseline method where sequenced sites were matched with the EPICv2 manifest prior to K-NN imputation using the impute.knn function (default parameters), 2) slideimp’s sliding window K-NN (k = 25), or 3) sliding window PCA (5 components) imputation, using 1,000-CpG windows with a 25-CpG overlap between windows. Imputation performance was evaluated by calculating the batch-adjusted correlation between the Horvath 2013 and PCHorvath1 epigenetic age estimates produced by each imputation method and the participants’ chronological age (Horvath, 2013; Higgins-Chen *et al*., 2022).

## Results

### Runtime and memory benchmarks on DNAm data

The runtime benchmarks (Figure 1B) show that slideimp’s K-NN imputation with 4 cores parallelization (green) is approximately 150x faster than methyLImp2 and 33x faster than imputePCA. In addition, slideimp’s PCA imputation optimizations result in 4.6x faster runtime than the original imputePCA function at ≈75,000 CpGs. Above ≈50,000 CpGs, slideimp’s Ball-tree K-NN imputation provides a 2x to 5x runtime reduction compared to the brute-force approach. Implementations in slideimp uses 30% less memory allocation compared to impute.knn and 10x to 100x less than methyLImp2 and imputePCA (Figure S1). On full microarray data, the grouping and targeted subset of CpG imputation features of slideimp’s K-NN imputation would result in more drastic runtime reductions.

### Accuracy of K-NN and PCA imputation vs. methyLImp2

We benchmark slideimp’s K-NN and PCA imputation accuracy against methyLImp2 in single tissue samples with increasing missing data rates. For these demographically diverse, blood derived DNAm data, K-NN and PCA imputation accuracy measures are comparable to methyLImp2 (Figure 1C). With no added missing data, PCA imputation performs best in terms of MAE while methyLImp2 shows better RMSE and RSQ. As we increase the missing data rate, methyLImp2’s performance declines more rapidly and can fail at a missingness rate ≥ 6%. This result is due to methyLImp2’s dependence on non-missing CpGs to impute missing patterns in other CpGs. Hence, sliding window K-NN and PCA imputation are more suitable for whole-genome DNAm data where nearly 100% of the sequenced CpGs in our EM-seq samples have some missing data.

### Accuracy of K-NN and PCA imputation grouped by chromosome vs. ungrouped

Next, we compare the accuracy of slideimp in multi-tissue samples where the imputations are grouped by chromosomes vs. ungrouped. In this data, PCA imputation expectedly outperforms K-NN in terms of accuracy (Figure 1D). Importantly, applying K-NN and PCA imputations grouped by chromosome is faster and more accurate than the respective ungrouped approaches.

### Epigenetics clocks analyses simulations under MCAR, MAR, or MNAR

Finally, we used Monte Carlo simulations to evaluate epigenetic clock regression analyses with K-NN and PCA imputation under different missing data mechanisms across increasing CpG missing rates. Figure S2 shows the coverage of effect size estimates. Estimates using PCA imputation are significantly less biased and have less variance than those of K-NN imputation under all missing data generating mechanisms, though both methods work well even up to 50% per-CpG missing rate. Figure S3 displays the coverage of the respective standard error estimates and PCA expectedly performs better than K-NN even at 20% per-CpG missing rate. Estimates from the K-NN Ball-tree methods (Figure S4, S5) are worse than using brute-force K-NN due to the required mean imputation prior to nearest neighbor identification. Overall, PCA imputation performs significantly better in all scenarios, but for lower per-CpG missing rates, K-NN provides satisfactory estimates. For both methods, the bias and variance of the estimators increase most under MNAR, followed by MAR, then MCAR, but are reliable at lower percentages of per-CpG missing rates.

### Calculation of epigenetic clocks from WGBS/EM-seq data with sliding window imputation

For the Horvath 2013 clock, slideimp increased the batch-adjusted correlation between chronological age and biological age from the baseline method of 0.131, p ≈ 0.382 to 0.325, p ≈ 0.039 for PCA imputation and 0.477, p ≈ 0.002 for K-NN imputation (Figure S6). PCClocks use information from 78,464 CpGs compared to the 353 Horvath 2013 clock CpGs and are more robust to measurement errors. slideimp improved the correlation between PCHorvath1 and chronological age from 0.735 to 0.797 for PCA imputation and 0.779 for K-NN imputation.

## Conclusion

slideimp provides the fastest K-NN and PCA imputation implementation in R for DNAm data from both microarray and whole-genome-based methods with high accuracy. Missing DNAm data imputation via slideimp’s sliding window imputation also facilitates the calculation of more accurate biological age estimates from whole-genome DNAm data than can be achieved using naïve K-NN imputation.

With declining per-sample costs of whole-genome DNAm sequencing, and the emergence of long-read sequencing approaches, we expect sample sizes of whole-genome DNAm datasets to increase, which will further improve the accuracy of sliding window imputation as implemented in slideimp. Finally, while we demonstrate the application of slideimp imputation using DNAm data, we expect this approach will improve imputation for a variety of high-dimensional datasets.

