## Supplemental Figures for "slideimp: Efficient Imputation for DNA Methylation Data"

S1

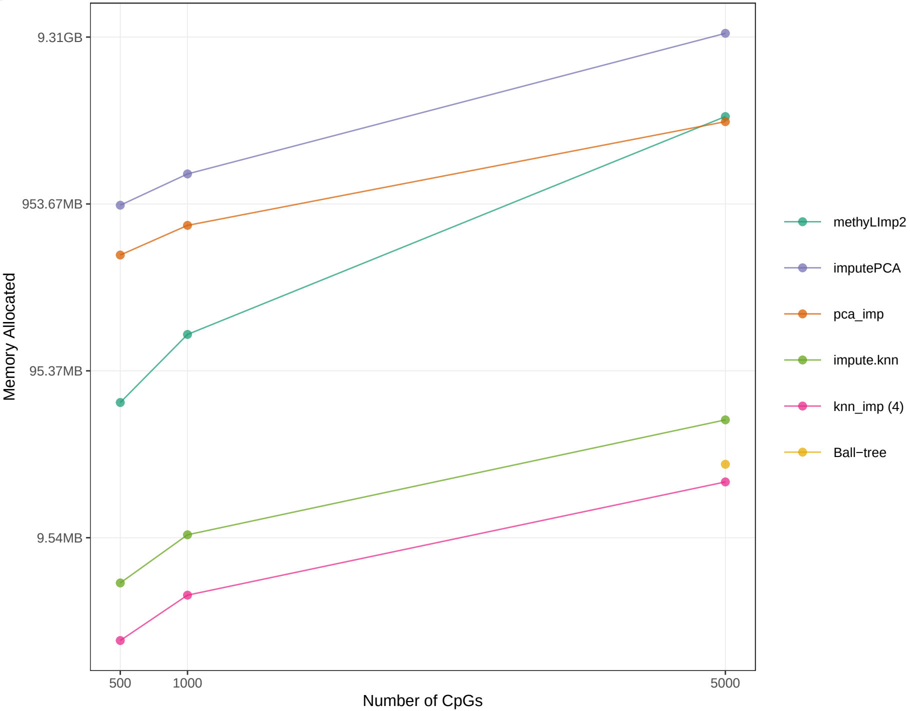


Figure S1. Memory allocation benchmarks for different imputation methods. CpG: cytosine–phosphate–guanine site.

**S2**

**Simulations, Effect Size Estimates Coverage, PCA and K−NN imputation**

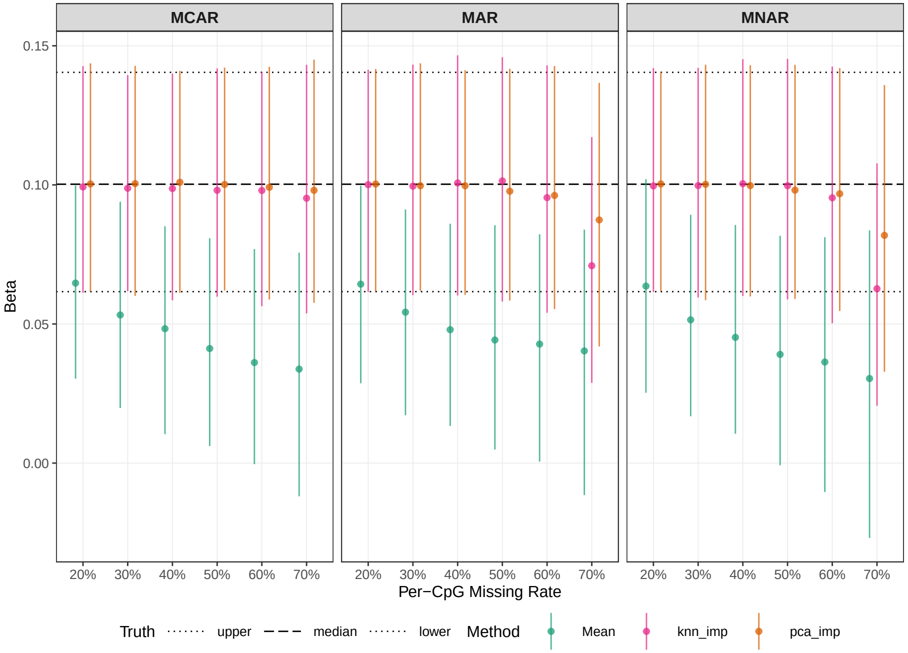


**S3**

**Simulations, Standard Error Estimates Coverage, PCA and K−NN imputation**

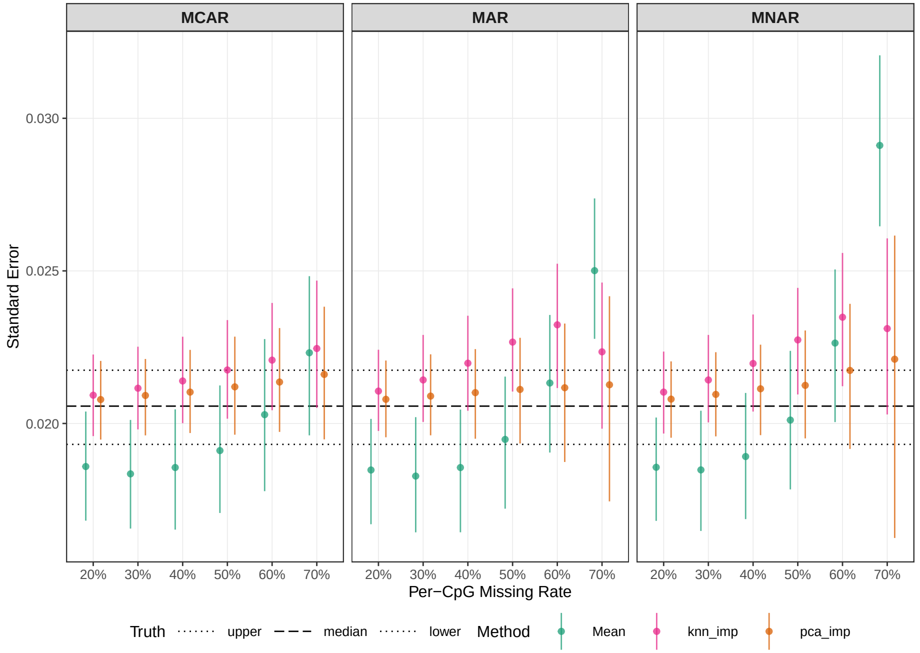


**S4**

**Simulations, Effect Size Estimates Coverage, Ball−tree K−NN imputation**

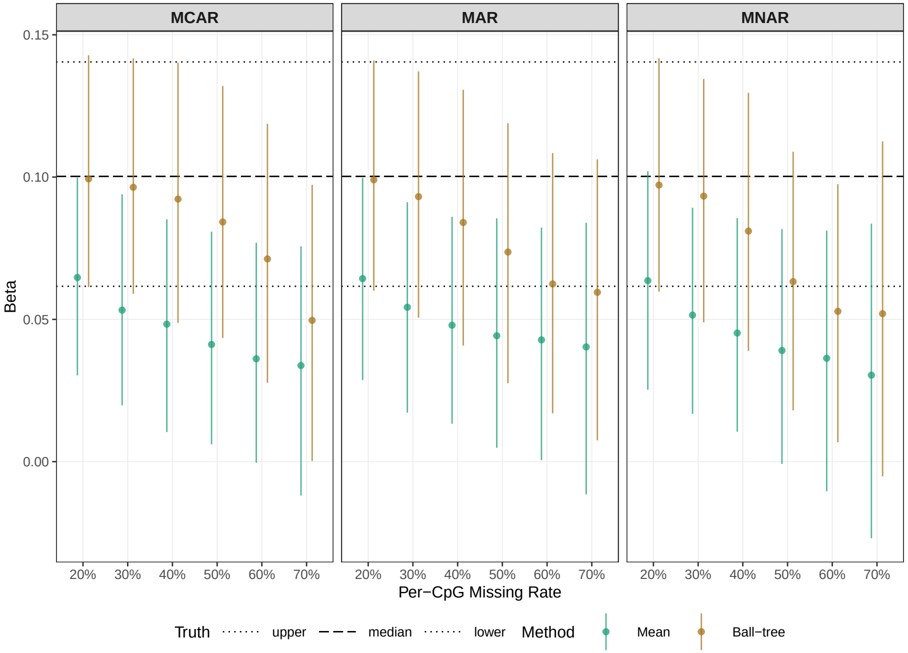


**S5**

**Simulations, Standard Error Estimates Coverage, Ball−tree K−NN imputation**

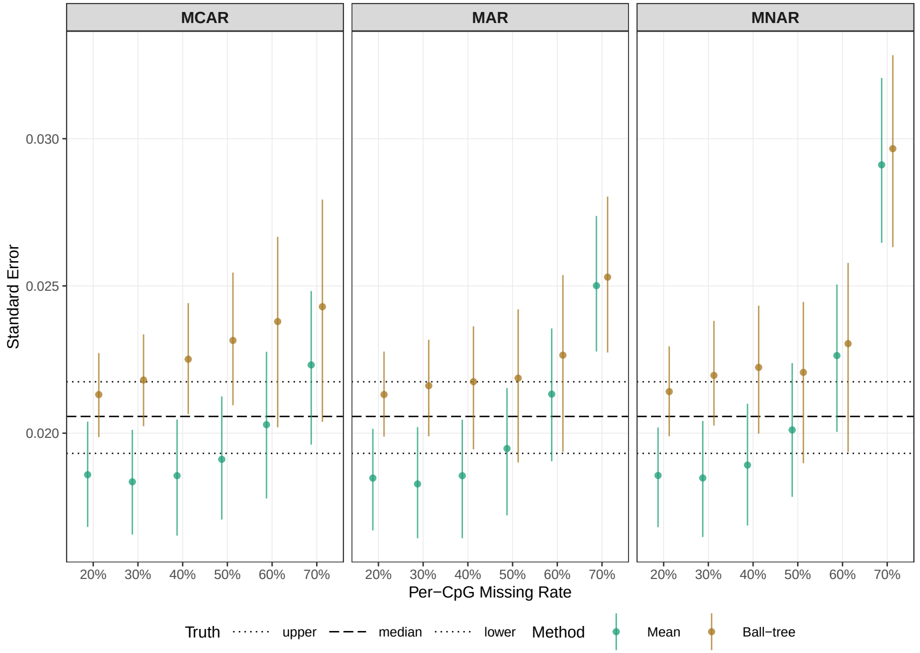


Figure S2, S3, S4, S5. Results from 1000 Monte Carlo simulations of regression estimates under Missing Completely at Random (MCAR), Missing at Random (MAR), and Missing Not at Random (MNAR) mechanisms, across different per−CpG missing rates. The points represent the medians of the distributions. The lines represent the 95% quantile intervals of the distributions. The dotted and dashed lines represent the known true values. PCA: principal component analysis; K−NN: k−nearest neighbors; CpG: cytosine−phosphate−guanine site.

S6

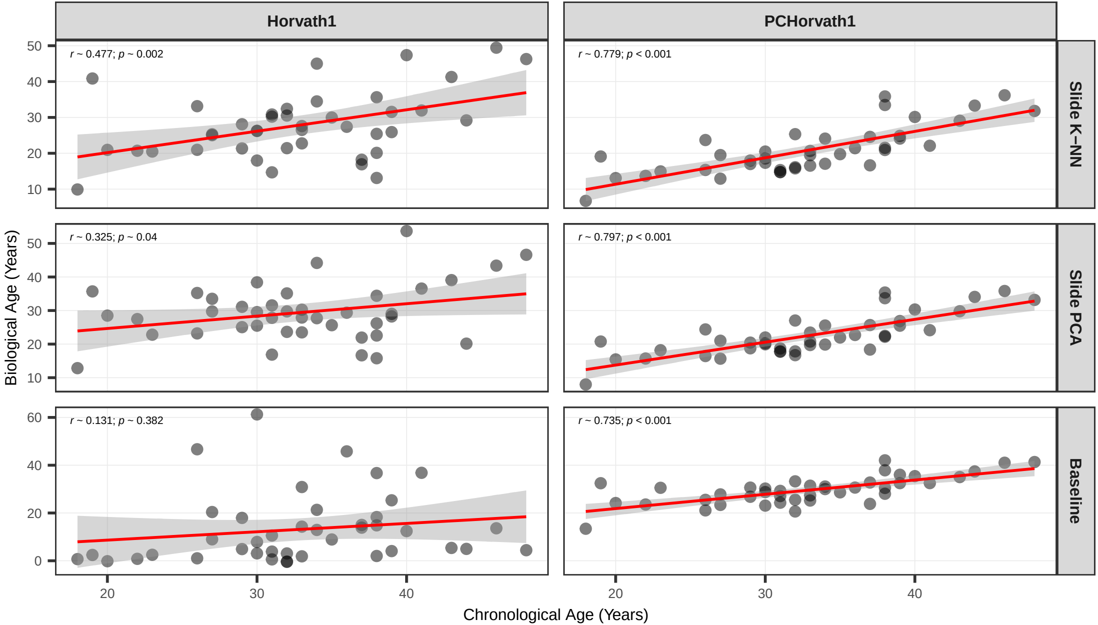


Figure S6. Estimation of epigenetic clocks using imputed EM−seq data from 41 reproductive−age women with no pregnancy or malignancy. The Horvath 2013 clock estimates are in the left column. The PCHorvath1 clock estimates are in the right column. From top to bottom, the imputation methods are sliding window K−NN, sliding window PCA, and the baseline method. Correlation values and p−values are calculated after adjusting for the batch of the samples. K−NN: k−nearest neighbors; PCA: principal component analysis.
